# Gene Interaction Network Architecture of Human Polygenic Traits Reveals Domain-Specific Connectivity and Evolutionary Pressures

**DOI:** 10.1101/2025.05.23.655809

**Authors:** Ehsan Tamandeh, Jessica Bigge, Adrian Serohijos, Johannes Schumacher, Carlo Maj, Pouria Dasmeh

## Abstract

Human polygenic phenotypes arise from complex interactions among genes within regulatory networks. To gain insights into the structural and evolutionary characteristics of these networks, we analyzed gene interaction networks across 4756 human polygenic phenotypes, contrasting the network properties of genes associated with polygenic phenotypes against those of non-associated genes. Our results indicate that genes associated with polygenic phenotypes exhibit a significantly higher connectivity in their corresponding gene-interaction networks, with connectivity varying markedly across different trait domains. Notably, genes within highly connected networks are enriched for immune-related biological processes, whereas genes in less connected networks are more involved in neurogenesis. Furthermore, we found that genes embedded in highly connected networks are, on average, under weaker selective constraints than those in lowly connected networks. Overall, our findings provide a systematic analysis of gene interaction networks underlying human polygenic traits and reveal how selective constraints vary with network connectivity. These insights offer a framework for prioritizing genes based on their connectivity within trait-associated networks. To support this effort, we developed the online portal http://netpolygen.com enabling researchers to generate and explore hypotheses by identifying genes that are both highly associated and highly connected across thousands of polygenic phenotypes.

## Introduction

Most human traits and diseases are polygenic, with multiple genes contributing to their manifestation through regulatory networks that facilitate gene-gene and protein-protein interactions^1–6^. Understanding these networks is crucial for gaining mechanistic insights into polygenic phenotypes and examining how genes associated with these traits may evolve differently from non-associated genes^4,7,8^. This is especially relevant for polygenic phenotypes, where the complex genotype-to-phenotype relationship and the distributed nature of selective signals across many genes make analyses focused on individual genes less informative.

Gene interaction networks are broadly categorized into physical and functional interaction networks^9^. Physical interaction networks built from protein-protein interactions map direct contacts between proteins, while functional networks connect genes based on co-expression, shared functions, or regulatory relationships. Both types of networks often form modular and hub-like structures, reflecting a common structural pattern in biological networks that likely enhances robustness against perturbations such as mutations and genetic aberrations^10–12^. While gene interaction networks have been elucidated in several polygenic phenotypes^2,13–17^, we know little about whether their structure varies across different traits. Furthermore, it remains unclear whether genes embedded within these networks are under unique selective constraints depending on their connectivity, potentially influencing their evolution.

To know more about gene interaction networks underlying human polygenic phenotypes, we systematically investigated the structure of these networks across 4756 traits from the GWAS ATLAS database^18^. We constructed interaction networks of highly associated genes with these phenotypes and contrasted them with the network of the least associated genes (see Methods). Our findings reveal that the interaction network of genes associated with polygenic phenotypes have a significantly higher connectivity compared to the network of non-associated genes. Furthermore, we found that the degree of connectivity varies across domains of polygenic traits—an effect not observed among non-associated genes. Immunological, skeletal, and metabolic traits exhibit the highest number of interactions per associated gene, while psychiatric traits show the lowest connectivity. Additionally, gene interaction network connectivity correlates with SNP heritability in polygenic traits: highly heritable traits have more connected gene interaction networks, with fewer non-interacting genes, compared to lowly heritable traits. Finally, we show that genes within highly connected networks of polygenic phenotypes are under weaker selective constraints compared to those in less connected networks. Notably, this pattern is observed only among associated genes, suggesting that phenotypic association, coupled with network connectivity, contributes to a relaxation of selective constraints specifically on these genes. Overall, our study provides a comprehensive view of the structure and properties of gene interaction networks underlying human polygenic phenotypes, illustrating how these networks may influence the selective pressures exerted on genes associated with these traits.

## Results

### Compilation of data and construction of gene interaction networks

To investigate the structure and properties of the interaction networks of genes associated with human polygenic phenotypes, we selected the available human polygenic traits in the GWAS ATLAS (4,756 traits, version database release 3, v2019; Figure 1A)^18^. We use the terms polygenic phenotypes and polygenic traits interchangeably throughout this work. GWAS ATLAS compiles this data by calculating the association of human genes (18,476 genes) with each of these traits using MAGMA, a statistical approach that aggregates the effects of SNPs and their association to the trait of interest for each gene. MAGMA assigns a *p*-value to every gene which represents its association with the trait of interest. We chose the genome-wide significance threshold of 2.7x10^-6^ (Bonferroni correction for the number or tested genes within MAGMA framework) and selected traits for whom at least 60 associated genes were present. This resulted in 461 traits (Tables S1-2). We applied this filtering step to reduce the impact of false-positive associations.

**Figure 1.**
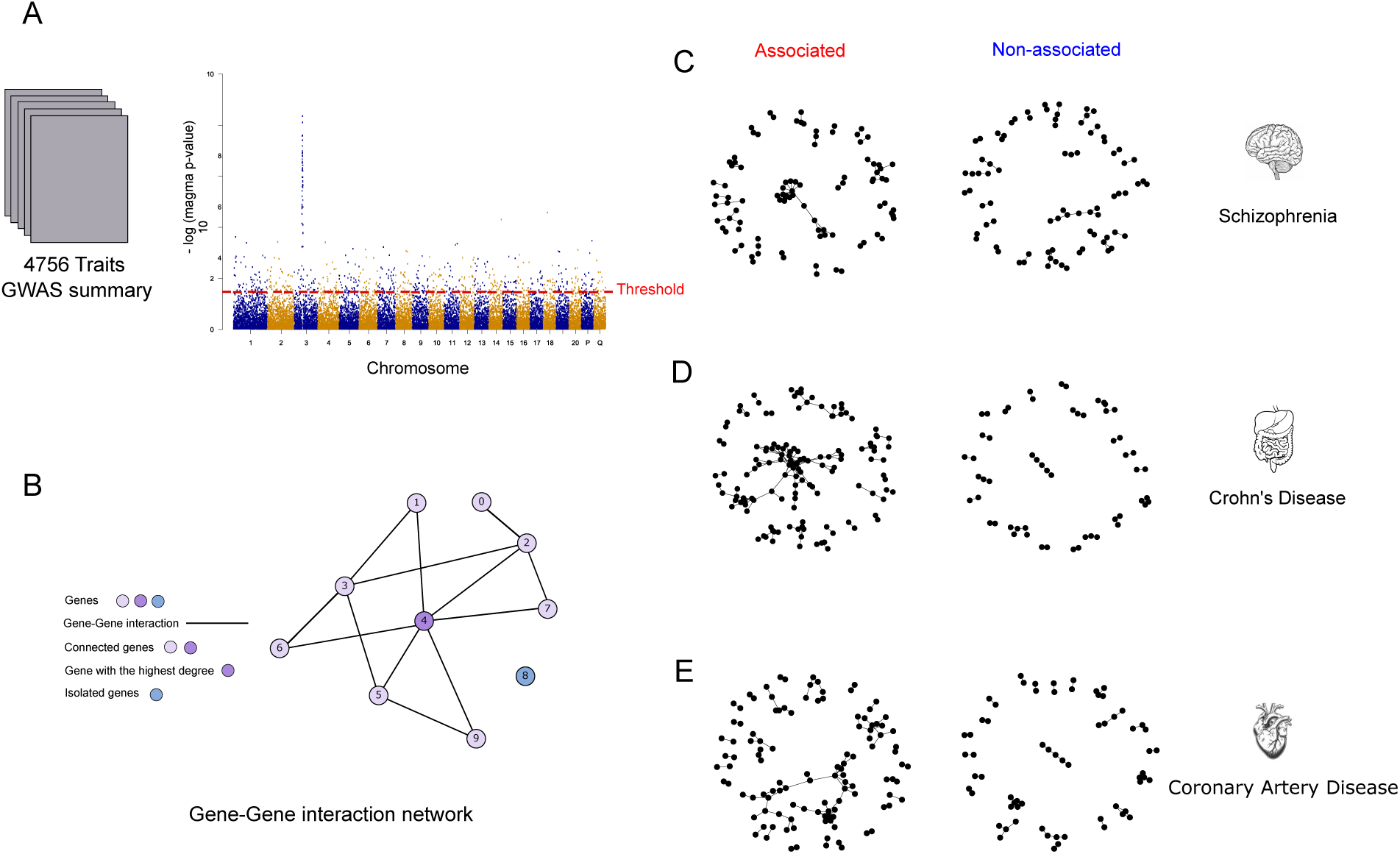
Gene-Interaction Network Analysis of Human Polygenic Phenotypes. A) An overview of our analysis. We identified significant gene–trait associations across 4,756 traits from the GWAS ATLAS and constructed gene interaction networks. To ensure robust network construction, we focused on traits with at least 60 significantly associated genes (p < 10⁻⁶). B) Schematic representation of a gene interaction network, highlighting connected genes, isolated genes, and genes with the highest connectivity. C-E) Gene interaction networks for associated and non-associated genes to schizophrenia (C, trait ID 9), Crohn’s disease (D, trait ID 68), and coronary artery disease (E, trait ID 108). This comparison illustrates the differences in network structure and genetic architecture between associated and non-associated gene networks for these polygenic diseases.

**Figure 2.**
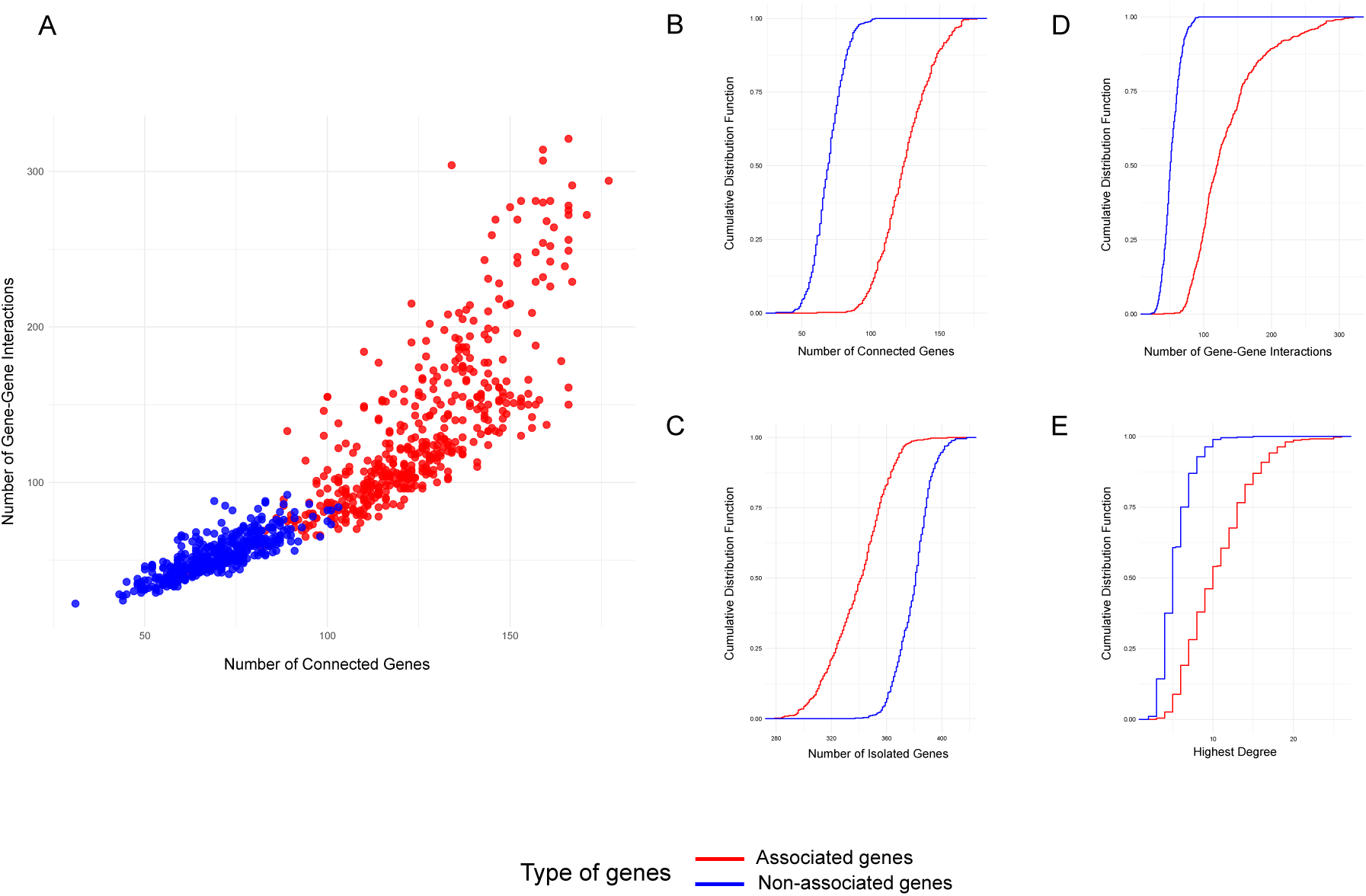
Comparison of Gene-Interaction Network Properties for Associated and Non-Associated Genes in Polygenic Phenotypes. A) The number of gene-gene interactions versus the number of connected genes in the interaction networks of associated genes (shown in red) and non-associated genes (shown in blue) with human polygenic phenotypes. Panels (B-E) present the cumulative distribution function (CDF) plots that compare various gene properties between associated and non-associated genes. In each plot, the y-axis represents the cumulative distribution, and the x-axis indicates the respective metric values: the number of connected genes (panel B), the number of isolated genes (panel C), the number of gene-gene interactions (panel D), and the highest degree of each network (panel D). The networks of associated genes (in red) and non-associated genes (in blue) were built using 500 highly, and 500 lowly associated genes, respectively. We selected the highly and lowly associated genes to each trait of interest from the MAGMA p-values. Comparisons of the network properties between associated and non-associated genes are highly significant, with *p* < 10⁻¹⁶ using the Wilcoxon rank-sum test.

However, when relevant, we also extended our analysis to the full set of 4,756 polygenic traits by relaxing this constraint.

We then retrieved interactions scores for these genes from the STRING database^19^ and constructed the gene interaction networks (see Methods for details) (Figure 1B, Figures S1-2). These networks were built using either physical or functional interaction scores, which we explicitly specify throughout this work when relevant. For either of these network types, we first constructed the “associated networks” from highly associated genes. For comparison, we constructed “non-associated networks” using an equivalent number of genes with the lowest association to the traits of interest.

We implemented two approaches to determine the number of genes used to construct associated and non-associated networks. In the first approach, we selected the top 500 most strongly associated genes and the bottom 500 least associated genes to build the associated and non-associated networks, respectively (see Methods). We chose this fixed number of genes for comparability across traits in our phenome-wide analyses, as this ensures uniformity in the statistical evaluation of network properties across diverse phenotypes. Selecting approximately 500-1000 genes is a common practice in polygenic analyses^20–22^ as it provides a balance between providing sufficient statistical power and capturing robust biological signals. In the second approach, we included only those genes meeting the genome-wide significance threshold (MAGMA *p*-value=2.7x10^-6^), matched with an equal number of the least-associated genes for each trait (see Figure S3 for details). We then analyzed the structure of these networks and examined their properties such as the number of edges, the number of connected and isolated nodes, and the degree distribution of each network.

### Genes associated with polygenic phenotypes have more interactions in gene networks than non-associated genes

Using the gene-gene interaction network of human polygenic phenotypes, we addressed the first question of our study: How do interaction networks differ between associated and non-associated genes? To answer this question, we compared the number of interactions and the network structure of associated and non-associated genes for three representative polygenic diseases: coronary artery disease, schizophrenia, and Crohn’s disease (Figures 1C-E). The genetic correlation between these traits is low (Rg ∼ 0.03) making them ideal candidates for our analysis because their network properties will be less likely affected from the same genes and interactions^7,23^. We then compared the interaction networks of 500 highly and 500 lowly associated genes with these polygenic diseases (see Methods for details). For all three traits, the interaction network of associated genes had a higher number of interactions and connected nodes, compared to the network of non-associated genes (Table 1). For example, in Schizophrenia, the network of associated genes had 86 interactions compared to 57 interactions that were present among the non-associated genes. This was also the case for coronary artery disease and Crohn’s diseases where the number of interactions between associated genes were 151 and 160 compared to 34 and 50 interactions among their non-associated genes, respectively. We also compared the degree distribution of both associated and non-associated networks to assess differences in network topology. We found that both networks followed a scale-free distribution, a characteristic commonly observed in biological networks (*p* < 0.05, Likelihood Ratio Test; Figure S4). This finding indicates that the primary difference between associated and non-associated networks lies in their connectivity rather than in their overall structure.

**Table 1.** Characteristics of Gene interaction Networks for Polygenic Phenotypes. This table provides a comparative analysis of gene interaction networks associated with schizophrenia (Trait ID 9), Crohn’s disease (Trait ID 68), and coronary artery disease (Trait ID 108). For each trait, we list the network type (associated or non-associated), the number of connected and isolated genes, the total count of gene-gene interactions, the maximum degree of connectivity among genes, and the slope of the degree distribution on a log-log plot.

| Traits | Network type | Connected genes (no.) | Isolated genes (no.) | Gene-gene interactions (no.) | Highest degree | Slope of degree distribution (log-log scales) |
| --- | --- | --- | --- | --- | --- | --- |
| Schizophrenia (Trait ID 9) | Associated | 94 | 369 | 86 | 11 | -1.74 |
|  | Non-associated | 81 | 376 | 57 | 4 | -2.77 |
| Crohn's Disease (Trait ID 68) | Associated | 139 | 315 | 151 | 13 | -1.79 |
|  | Non-associated | 57 | 392 | 34 | 3 | -3.38 |
| Coronary Artery Disease (Trait ID 108) | Associated | 115 | 355 | 100 | 8 | -1.97 |
|  | Non-associated | 65 | 388 | 50 | 5 | -2.29 |
| Average (for all 461 traits) | Associated | 124.92 ± 19.20 | 319.12 ± 21.40 | 133.38 ± 51.55 | 10.53 ± 4.17 | -1.88 |
|  | Non-associated | 69.37 ± 11.23 | 379.81 ± 12.80 | 52.63 ± 12.70 | 5.37 ± 1.91 | -2.22 |

We then aimed to extend our observations from the three representative polygenic phenotypes to a broader set of human polygenic phenotypes. We constructed the network of gene interactions from 500 highly and lowly associated genes for 461 traits (from 4756 traits within the GWAS ATLAS) for whom at least 60 associated genes were present (Table S2). Indeed, and for all these traits, the number of interactions among the associated genes was significantly higher than those for non-associated genes (*p* < 10^-16^, Wilcoxon rank-sum test).

Two factors may influence our observations on the differences between the interaction networks of associated and non-associated genes. First, the non-associated genes used in our comparisons may be shared across multiple polygenic phenotypes, potentially including genes with minimal impact on complex trait manifestation. Second, we compared networks of 500 genes with the highest and the lowest associations, which may affect our conclusions for traits with fewer associated genes. Including additional genes—especially those with associations falling below genome-wide significance— could alter network characteristics and impact our findings.

To address these concerns, we first assessed the pairwise overlap of associated genes across traits. We randomly selected 50 traits from our set of 461 well-powered GWAS traits, each with at least 60 associated genes. On average, the number of overlapping associated genes between trait pairs was approximately 51 ± 41 genes (11.07 ± 8.96% of the 500 associated genes), which was significantly higher than the overlap observed among non-associated genes (15 ± 13 genes; 3.37 ± 3.01%). This difference was statistically significant (Wilcoxon rank-sum test, p < 10⁻¹⁶). This finding aligns with previous studies showing that genes associated with polygenic traits are often pleiotropic^24,25^, where a single gene can influence multiple traits^13,18^. We then repeated the comparisons between associated and non-associated gene interaction networks, this time restricting the analysis to genes surpassing the genome-wide significance threshold (MAGMA *p*-value < 2.7x10^-6^). For this analysis, we selected 10 traits with varying counts of associated genes, covering different quantiles of the number of gene interactions (Table S4). The associated networks exhibited significantly higher connectivity (*p*= 0.00293, Wilcoxon rank-sum test). Altogether, these results demonstrate that highly associated genes with human polygenic phenotypes form interaction networks with markedly greater connectivity compared to non-associated genes.

### The connectivity of gene interaction networks varies in different domains of polygenic phenotypes

We now turn to the second question of our study: Does the average interaction degree of genes differ across domains of polygenic phenotypes? We hypothesized that gene connectivity may vary by trait domain, reflecting structural differences in gene interaction networks. Gene–gene interaction networks are known to exhibit a hub-like architecture^16,26^, in which a small number of genes have many interactions while most have few. Within this framework, it is plausible that genes contributing to different trait domains occupy distinct regions of the network, with varying levels of connectivity that may reflect differences in their regulatory or functional roles. To explore this hypothesis, we examined the number of gene interactions and the number of connected nodes within gene interaction networks in traits across different domains of polygenic phenotypes. Indeed, these properties varied significantly across different domains of polygenic phenotypes (ANOVA test, *p* < 2.2x10^-16^; Table S5).

Polygenic phenotypes within the domains skeletal, immunological, and respiratory phenotypes had the highest number of interactions, and the lowest number of non-interacting genes (isolated nodes) compared to other domains of polygenic traits (Figure 3A). Psychiatric traits, on the other hand, had the lowest number of gene interactions and the highest number of non-interacting genes (Figure 3C). We further compared these results with those obtained from non-associated genes within each domain of polygenic trait. In contrast to the gene interaction networks of associated genes, non-associated genes did not show a significant variation across the domains of polygenic traits, neither for the total number of interactions (ANOVA test, *p* = 0.21; Table S5), nor for the number of isolated nodes (ANOVA test, *p* = 0.66, Table S5) (Figures 3B-D). Altogether, these results demonstrate that the connectivity of gene interaction networks varies in different domains of human polygenic phenotypes

**Figure 3.**
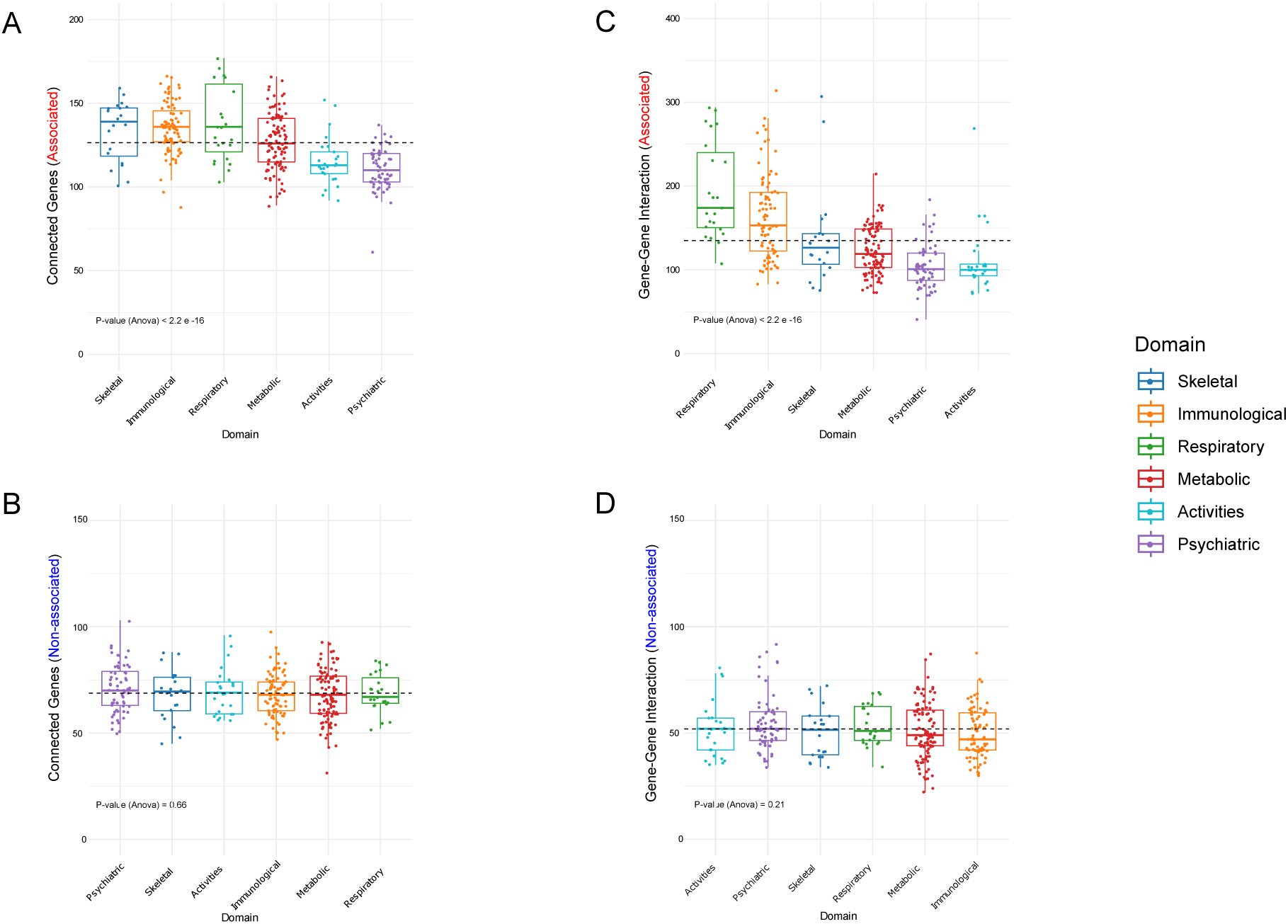
The connectivity of gene-interaction networks varies across different domains of polygenic phenotypes. Comparative analysis of gene-gene interactions and connected genes across different biological domains for associated and non-associated genes. Panels (A-D) represent gene interactions (A and B) and connectivity (C and D). A) Gene-gene interactions in associated genes across biological domains including Skeletal, Immunological, Respiratory, Metabolic, Activities, and Psychiatric. B) Gene-gene interactions in non-associated genes across the same domains. C) Number of connected genes in associated genes and, D) in non-associated genes across the biological domains. Statistical significance is indicated by ANOVA p-values, with a significant difference observed in Panels A and C (p-value < 2.2e-16), while panels B and D show non-significant variations (p-values of 0.66 and 0.21, respectively).

### The connectivity of gene interaction networks contributes to SNP heritability

In highly connected gene interaction networks, genes often participate across multiple pathways. Consequently, genetic variations affecting these genes might have a pleiotropic effect on several processes. This interconnectedness could contribute to variations in SNP heritability—the proportion of phenotypic variance explained by common genetic variants. A previous analysis of 42 complex diseases demonstrated that genes with high network connectivity contribute to SNP heritability^27^. To extend this result to the broader set of polygenic phenotypes, we analyzed SNP heritability of our study traits in relation to various network properties and assessed whether greater connectivity correlates with increased SNP heritability and, thereby, a stronger genetic impact on phenotypic variation. We found a significant positive correlation between SNP heritability (SNP h²) and the number of interactions (edges) within associated networks (*R*=0.29, p=3.93x10^-10^, Spearman’s rank correlation; Table S6), a relationship that was absent in non-associated networks (*R*=-0.06, p=0.10, Spearman’s rank correlation) (Figure 4A). To ensure that this result was not driven by the total number of genes in each network, we repeated the correlation analyses using partial correlations, controlling for the number of nodes. In these analyses too, SNP heritability showed a significant positive correlation with the number of interactions within associated networks (*R*=0.28, p=6.71x10⁻¹⁰, Spearman’s rank correlation), while no significant correlation was observed in non-associated networks (R=-0.05, p=0.28, Spearman’s rank correlation).

**Figure 4.**
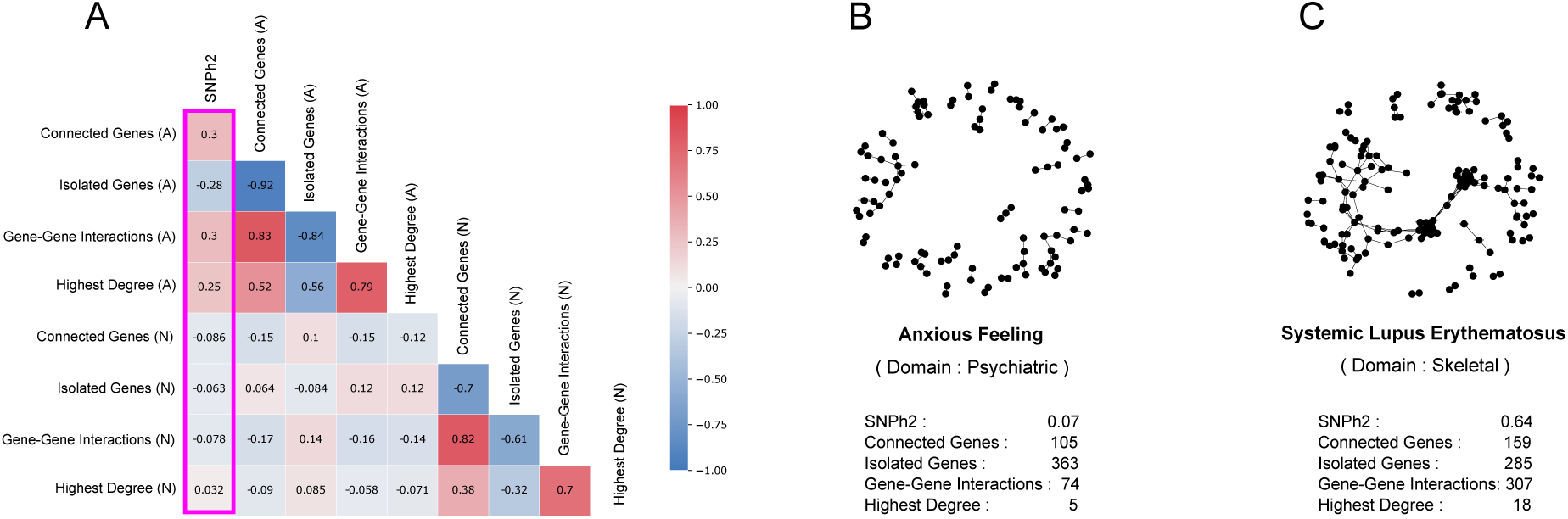
Contribution of Gene Interactions to SNP Heritability (h²) in Polygenic Phenotypes. A) The correlation diagram showing the Spearman’s rank correlation between SNP heritability of different polygenic traits and various properties of their gene interaction networks. B, C) The interaction network of genes associated with the polygenic phenotype of anxious feeling with SNP heritability (h^2^) of 0.07 (panel B), compared to the interaction network of genes associated with the polygenic phenotype of systemic lupus erythematosus with the SNP heritability (h^2^) of 0.64.

As an illustrative example, we selected two traits with gene interaction networks at the extremes of SNP heritability: ’anxious feeling’ and ’systemic lupus erythematosus,’ with SNP heritabilities of 0.07 and 0.64, respectively. For ’anxious feeling,’ a psychiatric phenotype with a low SNP heritability of 0.07, the interaction network contained 105 connected genes, 363 isolated genes, 74 gene-gene interactions, and a maximum degree of 5. By contrast, ’systemic lupus erythematosus,’ an autoimmune phenotype with a higher SNP heritability of 0.64, showed a gene interaction network with 159 connected genes, 285 isolated genes, 307 gene-gene interactions, and a highest degree of 18 (Figure 4B, 4C).

We also expanded our analysis to all 4,756 traits available in the GWAS Atlas— not just the subset of 461 traits studied with well-powered genome-wide association studies. For each trait, we constructed gene interaction networks using the 500 most highly associated genes, incorporating both physical and functional interactions. Across all polygenic phenotypes, we observed that the average and median number of edges, maximum degree, and total number of edges within each network show a significant correlation with SNP heritability (Table S8). Additionally, there were not biased by the GWAS power (Supplementary note 1, Table S10). Altogether, these findings show that polygenic phenotypes with a higher SNP heritability have underlying gene interaction networks characterized by a higher network connectivity.

### Selective constraints on genes associated with polygenic phenotypes vary with gene interaction network connectivity

We then turned to the third question of our study: could genes associated with polygenic phenotypes be subject to differential selection pressures that stem from their placement within gene interaction networks? This question arises because, traditionally, a relationship has been observed between network connectivity and evolutionary rate— genes with higher connectivity are under stronger selective constraints and tend to evolve more slowly^28,29^. However, our previous work and that of others suggests that genes associated with polygenic phenotypes may be subject to different selective pressures compared to non-associated genes with such traits or genes linked to monogenic diseases. For instance, we have shown that the degree of association between human genes and polygenic phenotypes exhibits significant heterogeneity in its correlation with evolutionary rate⁵. We therefore set out to investigate whether association with polygenic phenotypes modifies the established relationship between gene connectivity and evolutionary rate.

Our initial goal was to replicate the well-established observation that genes with a higher number of interactions tend to evolve under stronger selective constraint compared to genes with fewer interactions, independent of any trait association. We calculated the average interaction count (average degree in the network) for 15,062 human genes associated with polygenic traits across our subset of 461 traits studied with well-powered genome-wide association studies. This approach is akin to a mean-field approximation in statistical physics, where the effect of individual interactions is averaged out, allowing us to generalize the influence of connectivity on selective constraints. We correlated these average interaction counts with the probability of loss-of-function intolerance (pLI), a widely accepted measure of selective constraint in human genes^30^. The pLI metric estimates a gene’s intolerance to loss-of-function (LoF) mutations, such as protein-truncating variants, frameshifts, and essential splice-site mutations. A pLI score near 1 indicates a high intolerance to LoF mutations, while a score closes to 0 suggests tolerance, implying that such variants are either neutral or only weakly deleterious. Our findings confirmed that genes with a higher average interaction count tend to have lower pLI value (*R* = 0.17, *p* < 10^-16^, Spearman’s rank-sum test; Table S11), supporting the expectation that genes with more interactions are under stronger selective constraint. Importantly, this relationship is independent of trait association, as it reflects an average across all traits.

We then examined how our results might change when accounting for the association of human genes with polygenic phenotypes. Specifically, we tested whether a gene’s placement within an interaction network is linked to differences in the selective constraints acting upon it. To do so, we calculated the correlation between the average pLI of genes within gene interaction networks and the number of interactions in each network. Here, deviating from mean-field behavior, we observed that genes within highly connected networks—with greater numbers of interactions—displayed an even lower average pLI indicative of relaxed selective constraints (*R* = -0.46, *p* < 10^-16^, Spearman’s rank correlation; Figure 5A, Table S8). Interestingly, this correlation was exclusive to the interaction network of associated genes and did not hold for non-associated genes (*R* = 0.07, *p* = 0.09, Spearman’s rank correlation; Figure 5A, Table S8).

**Figure 5.**
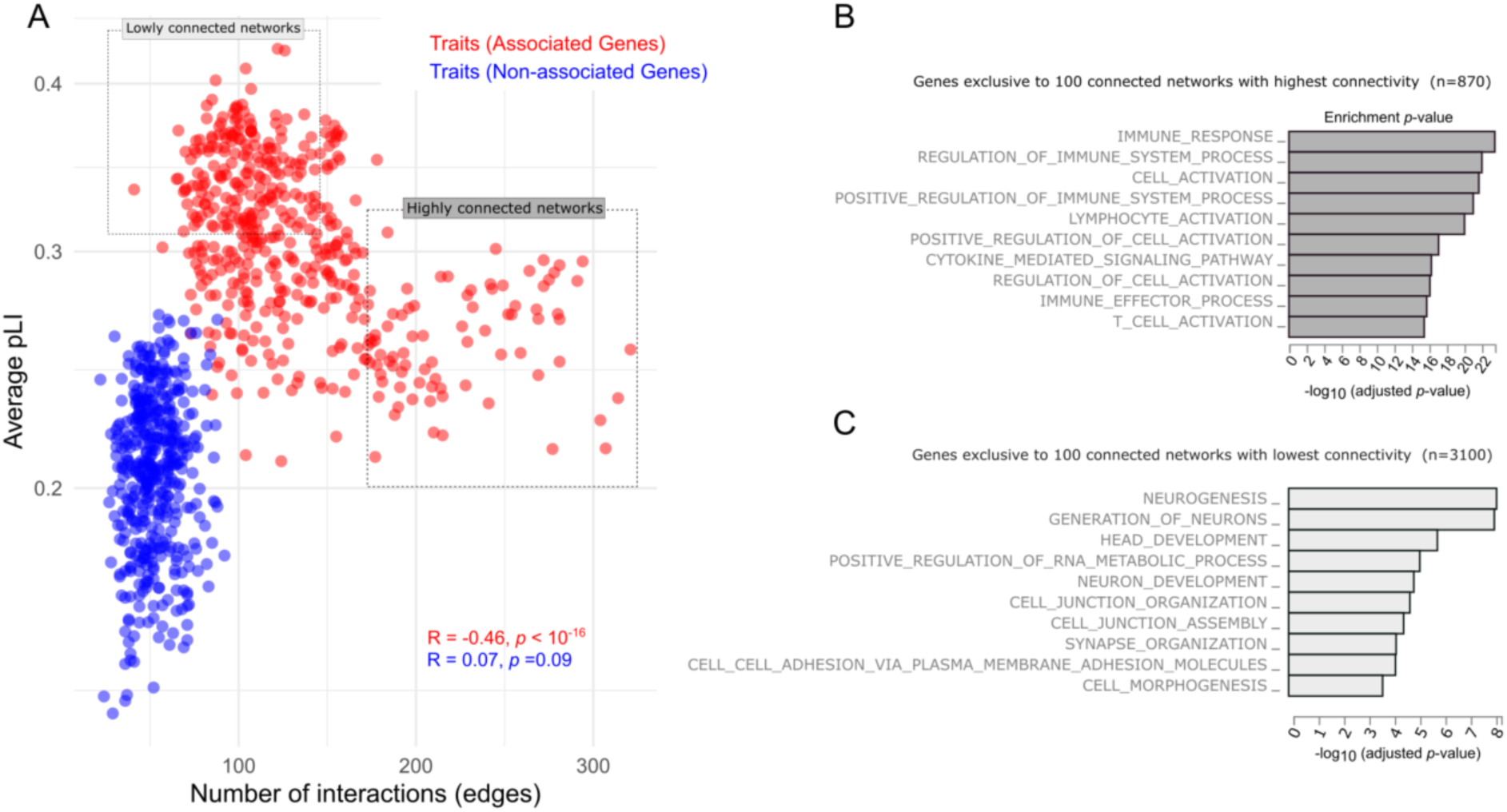
Selective Constraints on Genes Associated with Human Polygenic Traits. A) The average probability of loss-of-function intolerance (pLI) of genes within associated (in red) and non-associated (in blue) gene interaction networks plotted against the number of interactions (edges) in these networks. B, C) The enrichment of gene ontology (GO) terms related to biological processes within genes exclusive to 100 connected networks with the highest connectivity (panel B, n=870 genes), and 100 networks with the lowest connectivity (panel C, n=3100 genes). In both panels B and C, the x-axis shows -log10 of the adjusted p-value calculated from a hypergeometric test using Functional Mapping and Annotation of Genome-Wide Association Studies (FUMA).

We further repeated this analysis with another measure of selective constraint but focused on protein coding sequences. We calculated the correlation between network connectivity and the normalized ratio of the rate of nonsynonymous to synonymous mutations, d*N*/d*S*^31–34^. This ratio, d*N*/d*S*, is commonly used to assess the impact of natural selection on protein-coding sequences: values of d*N*/d*S* < 1 indicate purifying selection, while values of d*N*/d*S* = 1 or > 1 indicate neutral evolution or positive selection, respectively. Consistent with our results using pLI, we observed that genes embedded in highly connected gene interaction networks associated with polygenic phenotypes tend to have a higher average d*N*/d*S* (*R* = 0.23, *p* = 2.6 x 10^-7^, Spearman’s rank correlation; Figure S5). In contrast, there was no significant correlation between the average d*N*/d*S* of genes within non-associated networks and their number of interactions (*R* = 0.01, *p* = 0.79, Spearman’s rank correlation). Importantly, these correlations stayed significant even after correcting for the gene expression level which is the strongest determinant of d*N*/d*S* (Supplementary Note 2). Altogether, these results suggests that the evolutionary rate of genes associated with polygenic phenotypes is influenced by the structure of the gene-interaction networks in which they reside.

### Consistent Patterns of Gene Connectivity in Functional and Physical Networks

Thus far, our analyses have focused on functional interaction networks, which capture both direct and indirect relationships between genes, including co-expression, shared pathways, and regulatory associations. However, a key subset of gene–gene interactions arise from physical interactions between the protein products of genes—so-called protein–protein interactions (PPIs). These physical interactions are critical components of cellular function and important contributors to polygenic phenotypes^35^. To evaluate the robustness of our findings from functional networks, we asked whether similar patterns hold when restricting our analysis to physical interaction networks based solely on protein–protein interactions (see Methods). This approach enables us to assess whether the selective constraints and network properties observed in functional networks are also reflected in networks built exclusively from physical interactions.

We replicated our key findings from functional networks using protein–protein interaction (PPI) networks. First, genes associated with polygenic traits exhibited a significantly higher number of physical interactions compared to non-associated genes (Figure S8). This connectivity also significantly varied across different domains of polygenic phenotypes similar to the case of functional interactions (Figure S9). Second, traits whose associated networks contained more protein–protein interactions also tended to display higher SNP heritability (Figure S10). Third, associated genes with a greater number of physical interactions were, on average, under weaker selective constraint, as measured by both pLI and *dN/dS* metrics. Notably, this pattern was specific to associated networks and was not observed in networks of non-associated genes (Figure S11). Together, these results indicate that the structural and evolutionary signatures identified in functional networks are consistently reflected in physical interaction networks, reinforcing the robustness and generalizability of our conclusions.

## Discussion

Our findings reveal that gene interaction networks associated with polygenic traits exhibit a distinctive structure, characterized by higher connectivity compared to networks of non-associated genes. This connectivity varies across different domains of polygenic phenotypes, further underscoring the trait-specific genetic architecture of polygenic traits. Traditionally, polygenic architecture is understood mainly in terms of the number of genes associated with a trait^36,37^; however, our results highlight the importance of considering the interaction networks of these genes. Most importantly, as we demonstrate in this study, network structure significantly contributes to the SNP heritability of polygenic phenotypes, with the impact being most pronounced in highly connected networks.

We also observed that genes embedded in highly connected networks are, on average, under weaker selective constraint compared to those within less connected networks. It is important to clarify that this finding does not contradict prior observations that genes with a higher number of interacting partners generally evolve under stronger selective constraint^28,38,39^; indeed, our results confirm this pattern (Table S7). However, when focusing on associated genes with polygenic phenotypes and embedded in gene interaction networks, we find that these genes, on average, show patterns of relaxed selective constraints in highly connected networks relative to genes embedded in lowly connected networks.

To better understand the factors underlying the reduced selective constraints on genes embedded within highly interacting networks, we explored whether the functional characteristics of genes in highly and lowly connected networks contribute to these differences. We selected genes exclusive to 100 traits with the highest connectivity (n=870 genes) and 100 traits with the lowest connectivity (n=3100 genes), respectively. We applied a statistical procedure to identify the optimal threshold that maximizes the separation between highly and lowly connected traits. The most pronounced separation was observed at a threshold of 160 edges (see Methods for details, Figure S7). We then calculated the enrichment of biological processes within these gene sets (see Methods for details). Genes exclusive to highly connected networks were enriched in terms related to immune response, immune system processes, and immune cell activation, including lymphocyte activation and T cell activation (Figure 5B). HLA and important players in immune response (IL, STAT, HLA) are genes are within the highly connected genes, local Network clusters of JAK-STAT (p=9.02x10^-9^) and interleukin (p=3.3x10^-7^). Conversely, genes exclusive to lowly connected networks were enriched in processes such as neurogenesis, neuron development, and synapse organization (Figure 5C).

The overrepresentation of immune-related genes among those exclusive to highly connected networks suggests that functional redundancy may underlie the observed relaxation of purifying selection in these networks. Functional redundancy is a well-established feature of the immune system. It enhances the robustness of the immune system, allowing it to tolerate loss-of-function mutations without significant functional disruption^40^. For example, loss-of-function mutations in pattern-recognition receptors (PRRs) seldom result in infection susceptibility, as compensatory pathways mitigate their effects^41^. Similarly, alleles of *MBL2*, encoding mannose-binding lectin (MBL), which activates the lectin complement pathway, are common globally (frequencies up to 30%^42,43^) despite being linked to increased infection risk^44,45^. Alternative mechanisms, such as ficolins or the C1q-dependent classical pathway, compensate for MBL deficiency, illustrating the robustness of the immune system^46,47^. This robustness of immune-related phenotypes may contribute to the functional redundancy of the associated genes and underlies their relaxed selective constraints acting on them. Understanding how such network-mediated relaxation of selective constraints may have contributed to evolutionary adaptation is an intriguing question for future research.

Gene interaction networks further enable us to move beyond linear genomic annotations. Genetic variants often act through epistatic interactions or influence phenotypes via shared biological pathways—relationships that traditional fine mapping approaches may overlook while network-based analyses account for these complex interaction patterns. To provide an illustrative example, we plotted the gene interaction network for coronary artery disease (CAD) (Figure 6). Notably, connectivity within this network was not evenly distributed; a subset of hub genes, including APOE (encoding Apolipoprotein E), SMAD3 (encoding a key signal transducer for transforming growth factor beta receptors), and LPL (encoding lipoprotein lipase) are highly associated genes with the largest number of interactions in this network. These hub genes are well-established players in lipid metabolism and inflammation—key pathways in CAD pathogenesis^48,49^. This example demonstrates that prioritizing genes based on their network connectivity and degree of interaction, particularly in highly connected nodes, could reveal genes with substantial impact on phenotypic variance and trait heritability.

**Figure 6.**
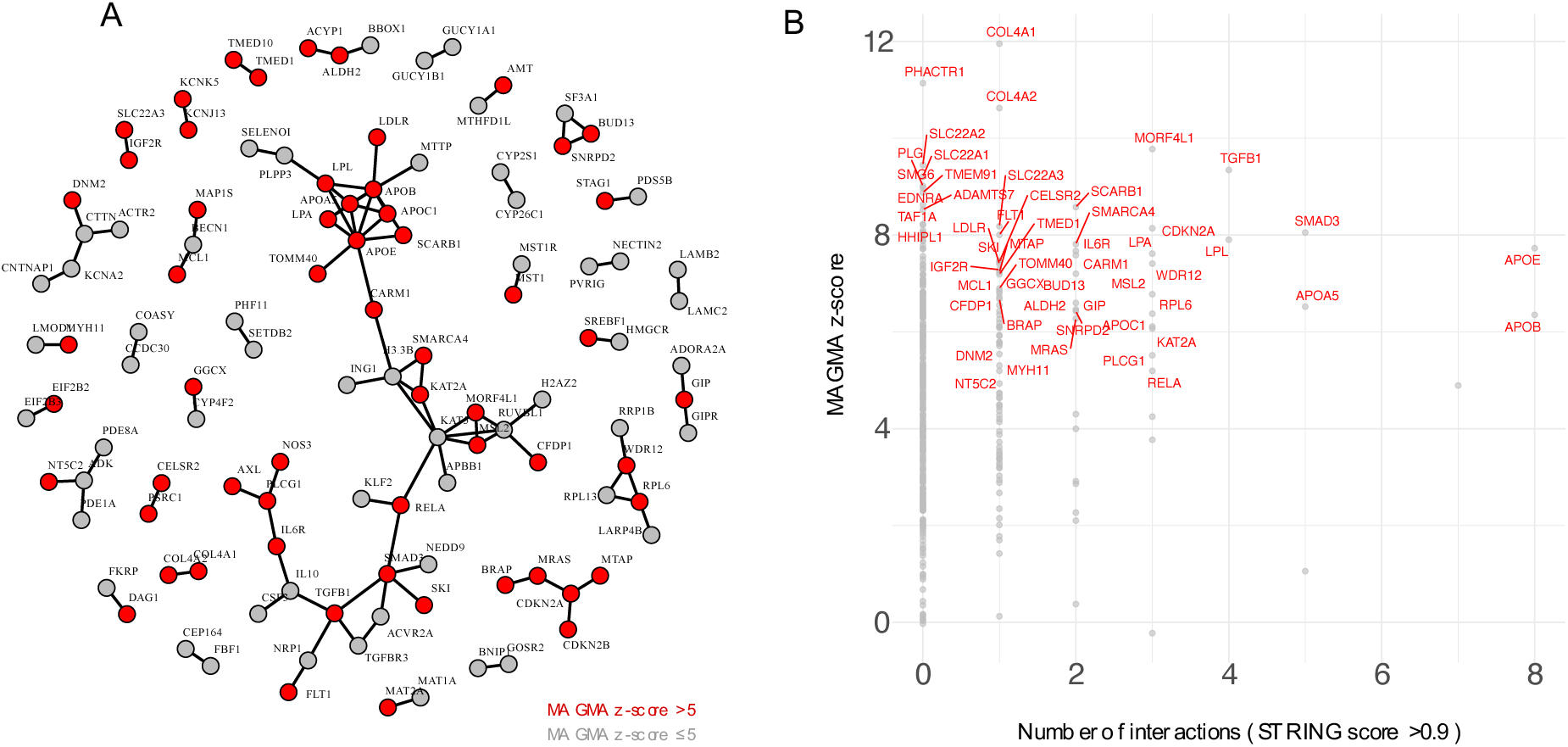
Gene-gene interaction networks facilitate the prioritization of GWAS-identified genes. A) The interaction network of genes associated with coronary artery disease (Trait ID 108 in the GWAS Atlas). Genes with a MAGMA z-score > 5 are highlighted in red. B) Scatter plot of MAGMA z-scores against the number of gene interactions for coronary artery disease-associated genes. Genes with a MAGMA z-score > 5 are labeled in red. This figure demonstrates that the number of interactions varies among highly associated genes, providing a framework for gene prioritization in fine-mapping analyses. For example, APOE (encoding Apolipoprotein E), SMAD3 (encoding a key signal transducer for transforming growth factor beta receptors), and LPL (encoding lipoprotein lipase) are highly associated genes with the largest number of interactions in this network.

Lastly and to facilitate hypothesis generation in this field, we developed NetPolyGen, a publicly accessible web portal that presents the network structure and properties of gene interaction networks associated with human polygenic traits. By integrating network-based insights, NetPolyGen enables researchers to identify highly connected, trait-associated genes and prioritize those most likely to be functionally significant. This resource aims to advance the study of complex trait biology by providing a systems-level framework for understanding genetic architecture and guiding future experimental and evolutionary investigations.

## Methods

To analyze the structure and topology of gene-gene interaction networks for human traits, we used the GWAS ATLAS database (version database release 3, v2019), which includes 4,756 polygenic traits and associations for 20,188 genes. Using MAGMA, a statistical tool that aggregates SNP effects to evaluate gene-trait associations, we calculated a z-score and corresponding p-value for each gene in relation to each trait. To ensure robust associations, we applied a genome-wide significance threshold of *p* < 10^−6^ and focused on traits with at least 60 associated genes, reducing our analysis to 461 traits (see Table S1).

For each of these traits, we constructed gene interaction networks using data from the STRING database (version 12.0, 2023), combining physical and functional interaction scores with high confidence (confidence score > 0.9). These networks, composed of genes with high association scores (termed "associated networks"), were then compared with networks constructed from the same number of genes with the lowest association to each trait (termed "non-associated networks"). To explore the structure of these networks, we analyzed properties such as the number of edges, the quantity of connected and isolated nodes, and the distribution of interaction degrees across nodes, allowing us to characterize each network’s complexity and connectivity.

We examined the degree distribution of each network by fitting three types of distributions: a power-law distribution, which corresponds to the degree distribution of scale-free networks; a Gaussian distribution, which represents the random network structure of the Erdos-Renyi model; and a log-normal distribution, often observed in biological networks with heavy-tailed distributions, such as metabolic networks. To assess the best-fitting model, we compared the likelihood of each fit using the Akaike Information Criterion (AIC).

To compare the gene ontology and functional enrichments of genes exclusive to highly and lowly connected networks, we applied a statistical procedure based on the Wilcoxon rank-sum test across a range of edge thresholds (Figure S7). Specifically, we tested whether gene constraint metrics, such as the average pLI score, differed significantly between genes in networks with low versus high connectivity. For each threshold, we calculated the p-value of the Wilcoxon test comparing the distributions of average pLI scores between networks with fewer versus more edges than the threshold. The strongest separation was observed at a threshold of 160 edges, which yielded a highly significant *p*-value of 9.32 × 10^-26^, indicating that threshold selection plays a critical role in shaping network-based inferences.

We used Functional Mapping and Annotation of Genome-Wide Association Studies (FUMA)^22^ to calculate Gene Ontology (GO) enrichment for genes associated with different GO terms, selecting key biological processes as the most insightful terms. Protein-coding genes served as the background set, and gene expression data from 30 general tissue types in the GTEx database (version 8) were used for the expression analysis. To account for multiple testing, we applied the Benjamini-Hochberg false discovery rate (FDR) correction, setting a maximum adjusted *p*-value threshold of 0.05 and requiring a minimum overlap of 2 genes with the gene sets.

We applied the gene-analysis module from the MAGMA tool to identify gene-association for coronary artery disease^50^. MAGMA computes *p*-values per gene by aggregating SNP-level association statistics within or near each gene to create a gene-level test statistic. An empirical null distribution is generated through permutations or simulations to represent the expected distribution of the test statistic under the null hypothesis. The p-value for each gene is then calculated as the proportion of null distribution values that are equal to or higher than the observed test statistic, indicating the gene’s statistical significance in association with the trait or disease of interest. In order to account for correlations between nearby genetic variants, MAGMA incorporates linkage disequilibrium structure of the regions. In our analysis, we accounted for linkage disequilibrium structure using the European LD matrix from Phase 3 of 1000 Genomes as provide by MAGMA (https://ctg.cncr.nl/software/magma). In order to compute gene-based association for CAD the genome-wide association study (GWAS) summary statistics from the Cardiogram consortium were used (available in: http://www.cardiogramplusc4d.org/media/cardiogramplusc4d-consortium/data-downloads/cad.additive.Oct2015.pub.zip)^51^.

All analyses were conducted in R (v4.3.3). The scripts to query the STRING database and construct networks are available in our GitHub repository: https://github.com/EhsanTamandeh/PolyGenNet-. The adjacency matrices for the interaction networks built from functional and physical interactions are available for download at our webserver: www.netpolygen.com

