## Supplementary Information for "Gene Interaction Network Architecture of Human Polygenic Traits Reveals Domain-Specific Connectivity and Evolutionary Pressures"

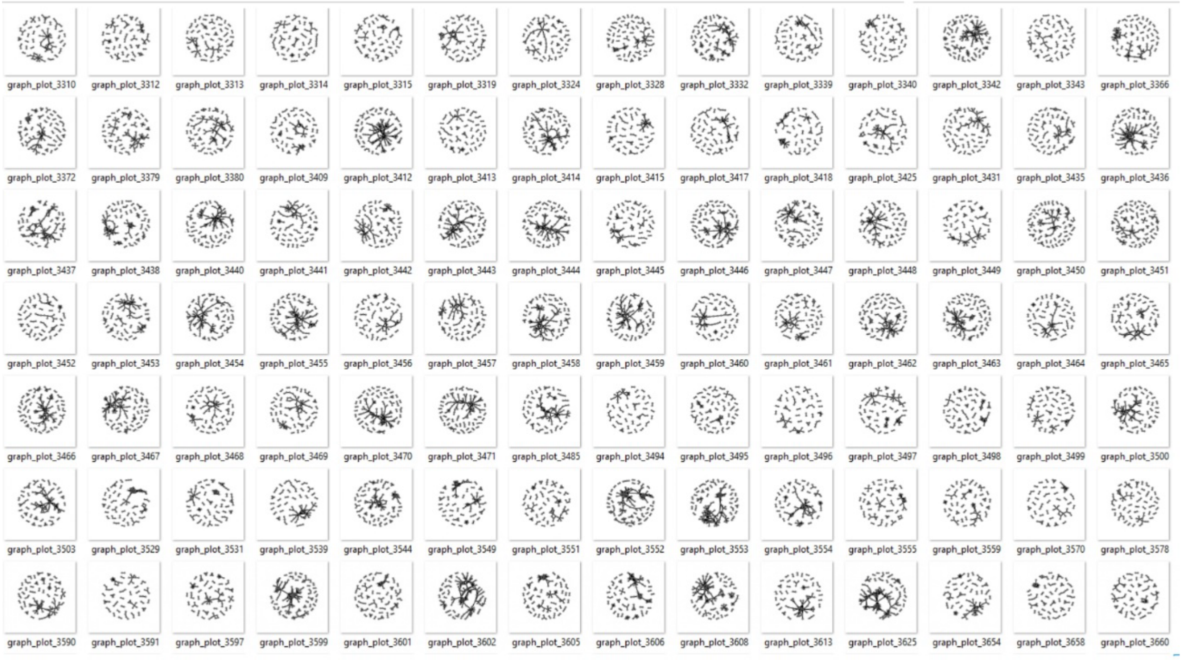


**Figure S1. The snapshot of selected associated networks.** The figure shows various structures of associated gene-gene interaction networks, illustrating the diversity in connectivity patterns. These networks exhibit a higher number of nodes and edges, indicating more gene-gene interactions and an increased presence of hub nodes, which play a central role in network connectivity. The presence of these hubs suggests a scale-free structure in the networks, highlighting the importance of certain genes in regulating the interaction dynamics.

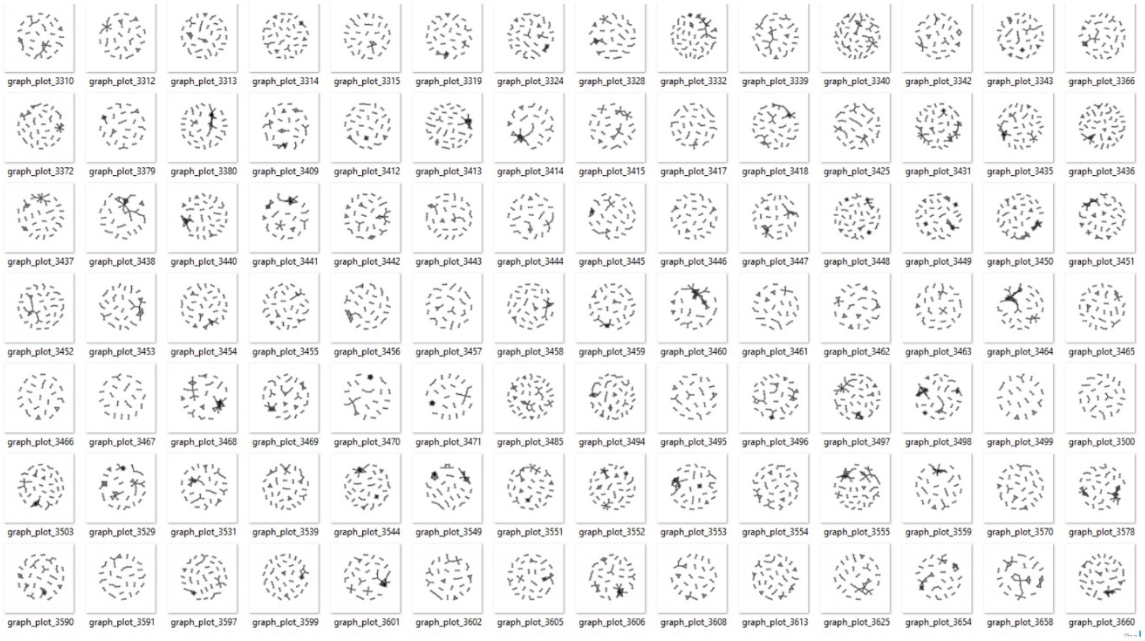


**Figure S2. Snapshot of selected non-associated networks.** This figure displays the structure of various non-associated gene-gene interaction networks, illustrating distinct patterns of connectivity with fewer nodes, edges, and hubs compared to associated networks (Figure S1). The non-associated networks demonstrate a lower degree of connectivity, reflecting limited interaction among nodes and a lack of dominant hubs that typically characterize highly connected networks.

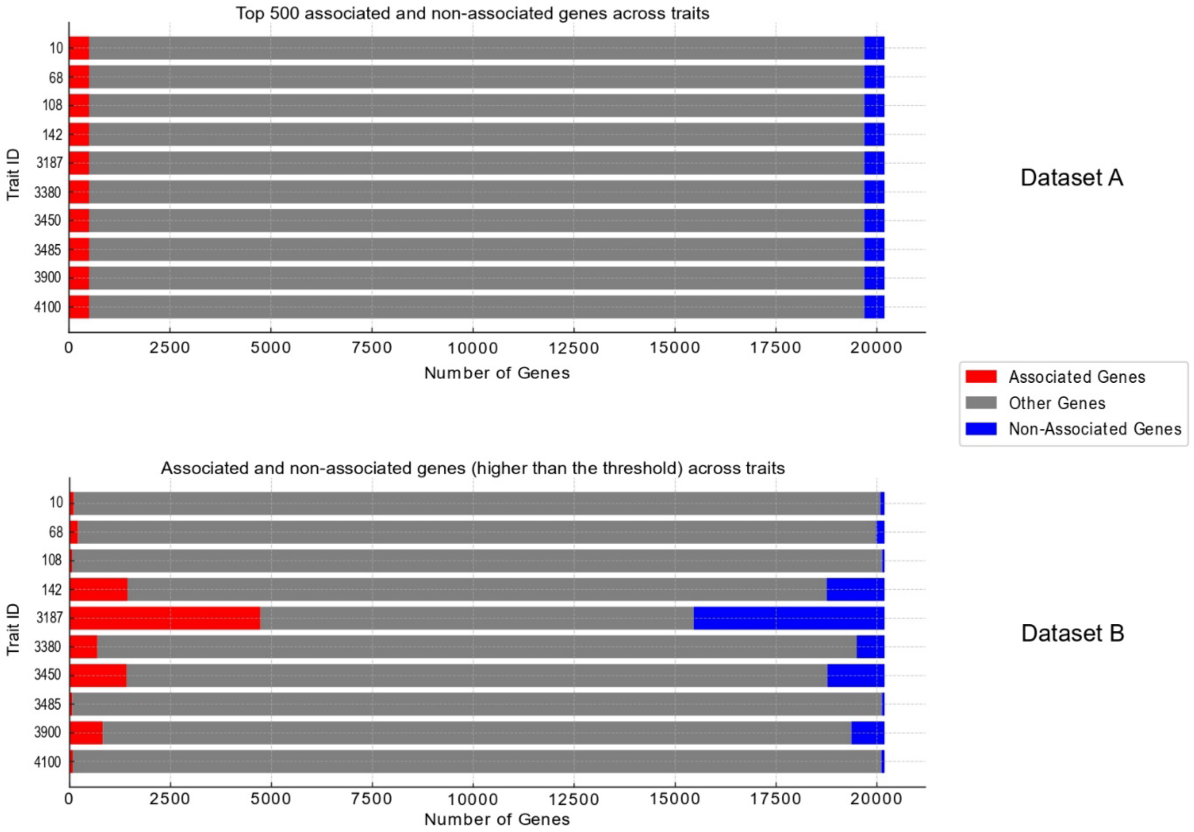


**Figure S3. Two different approaches to select associated and non-associated genes.** This figure compares gene distributions across various traits for two datasets, visualizing different selection criteria. On the upper panel, both associated and non-associated genes are displayed, with the top 500 genes shown for each trait, regardless of association strength. This provides a broader view of gene presence across traits. In contrast, the lower panel focuses exclusively on associated genes that exceed a specific threshold, offering a more targeted perspective on high-association genes across traits.

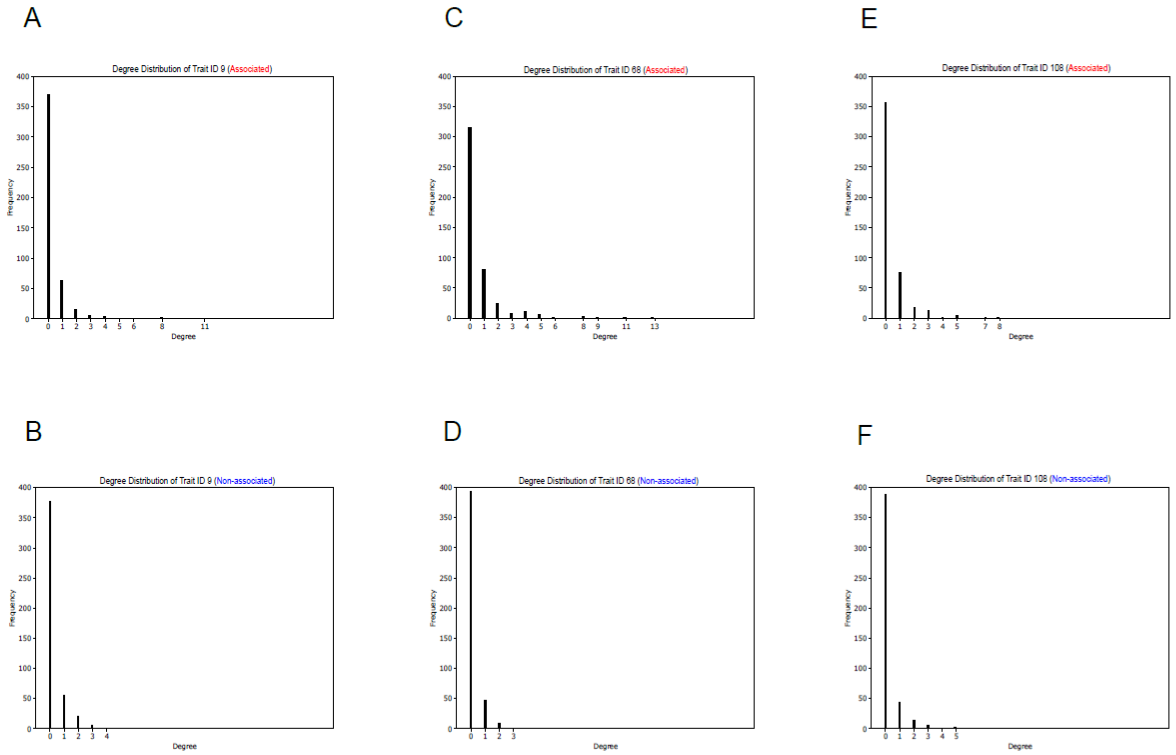


**Figure. S4**. Degree distribution of associated (panel A, C, E) and non-associated (panel B, D, F) networks in Schizophrenia, coronary artery disease, and Crohn's Disease. The x-axis represents node degree, while the y-axis indicates the frequency of nodes with each degree. These power-law distributions highlight a higher frequency of isolated nodes in non-associated networks and with higher number of nodes in associated network types, with higher number of interactions, showing higher connectivity, reflecting hubs and dense interaction patterns common in these networks. The presence of these distributions suggests a scale-free structure in the both network types. Degree Distribution of Networks for Different Trait IDs. (A, B) Degree distribution for Trait ID 9 (Schizophrenia): The associated network (A) exhibits a broader degree distribution, with nodes having degrees up to 11, whereas the non-associated network (B) has a more restricted range, with most nodes having degrees ≤ 4. (C, D) Degree distribution for Trait ID 68 (Crohn's Disease): The associated network (C) contains nodes with degrees up to 13, showing a wider spread compared to the non-associated network (D), where most nodes have degrees ≤ 3. (E, F) Degree distribution for Trait ID 108 (coronary artery disease): Similar to previous cases, the associated network (E) has a broader degree distribution (max degree 8) compared to the non-associated network (F), where node degrees do not exceed 5.

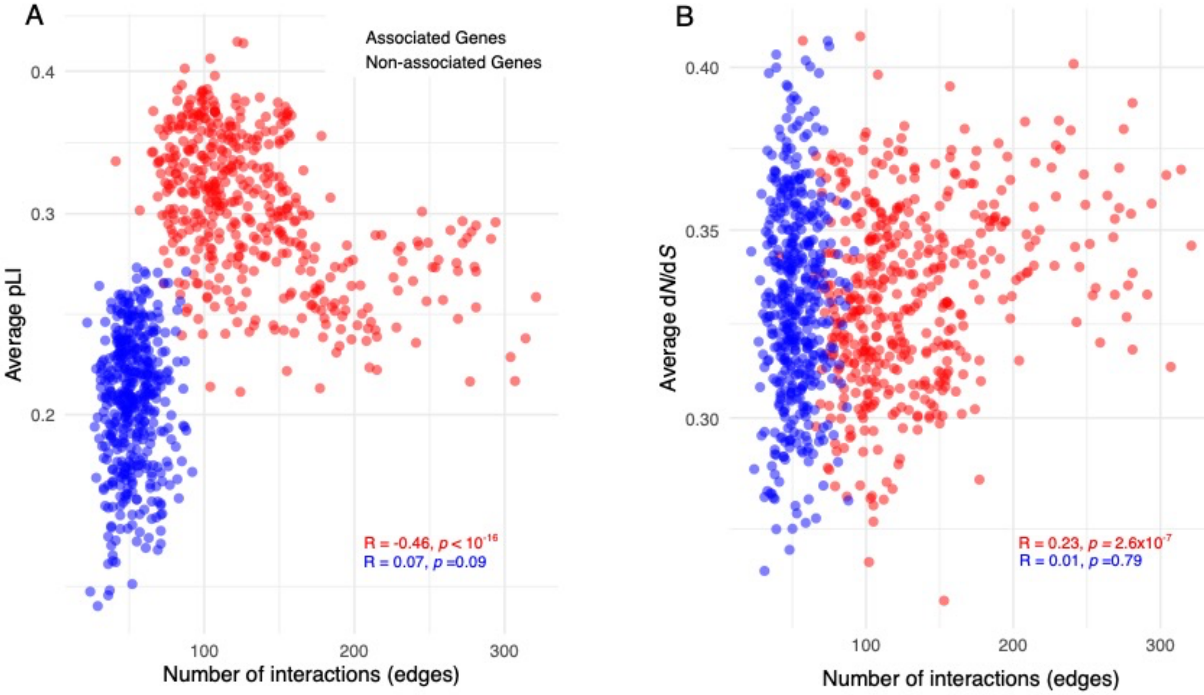


**Figure S5**. A) The average probability of loss-of-function intolerance (pLI) of genes within associated (in red) and non-associated (in blue) gene interaction networks plotted against the number of interactions (edges) in these networks. B) The average ratio of non-synonymous to synonymous substitution rates (dN/dS) for genes within associated (in red) and non-associated (in blue) gene interaction networks plotted against the number of interactions (edges) in these networks. In both panels, Spearman's correlation coefficient is used to assess the correlation.

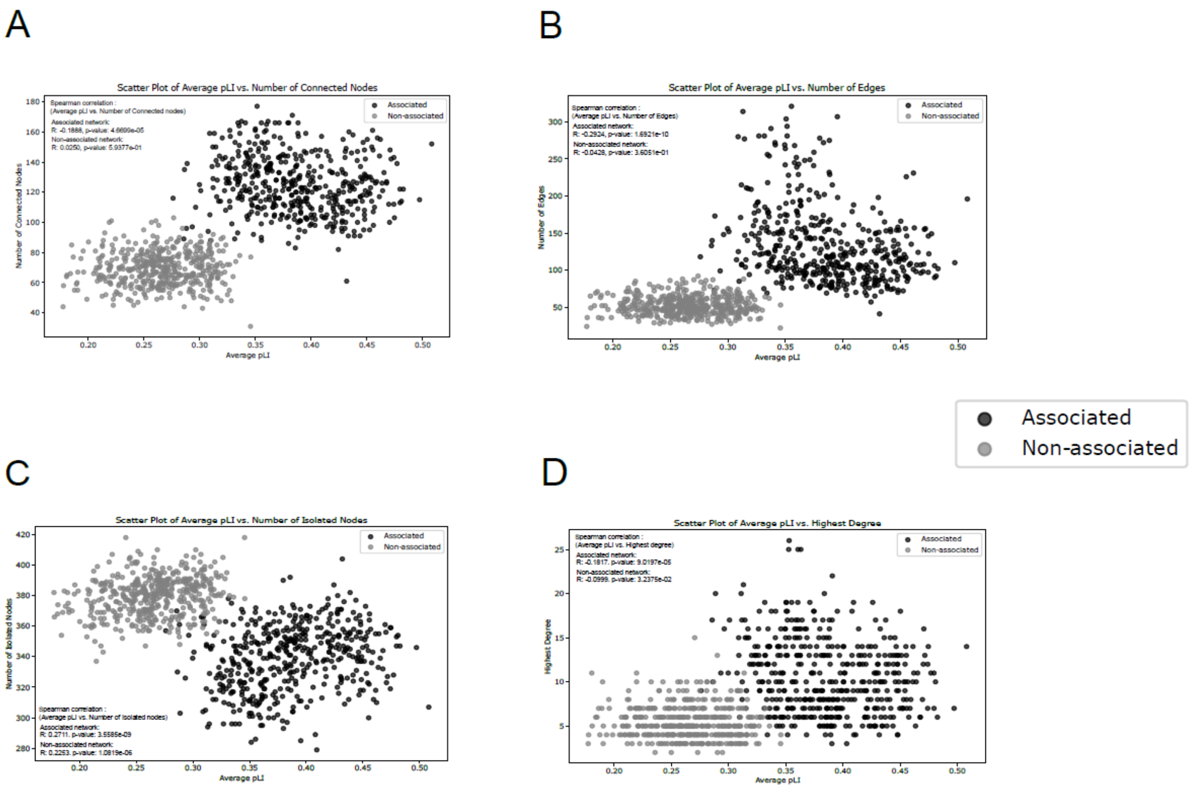


**Figure S6**. A) The average pLI vs. the number of connected nodes in networks built from functional scores: In the associated network, a weak negative correlation (R = -0.1888, p = 4.67e-05) suggests that highly connected genes tend to have lower pLI scores. No significant trend is observed in the non-associated network (R = 0.0250, p = 0.59). B) The average pLI for genes within functional networks vs. the number of edges: a moderate negative correlation is observed in the associated network (R = -0.2924, p = 1.69e-10), while the non-associated network shows no significant trend (R = -0.0428, p = 0.36). C) The average pLI vs. the number of isolated nodes: Both networks show a positive correlation, with isolated genes tending to have higher pLI values (associated: R = 0.2711, p = 3.56e-09; non-associated: R = 0.2253, p = 1.08e-06). D) The average pLI of genes vs. the highest degree in the network: a negative correlation suggests that highly connected genes tend to have lower pLI scores (associated: R = -0.1817, p = 9.02e-05; non-associated: R = -0.0999, p = 0.032). These findings suggest that in the associated network, higher connectivity is linked to lower gene constraint, while isolated genes exhibit greater functional constraint. This trend is weaker in the non-associated network.

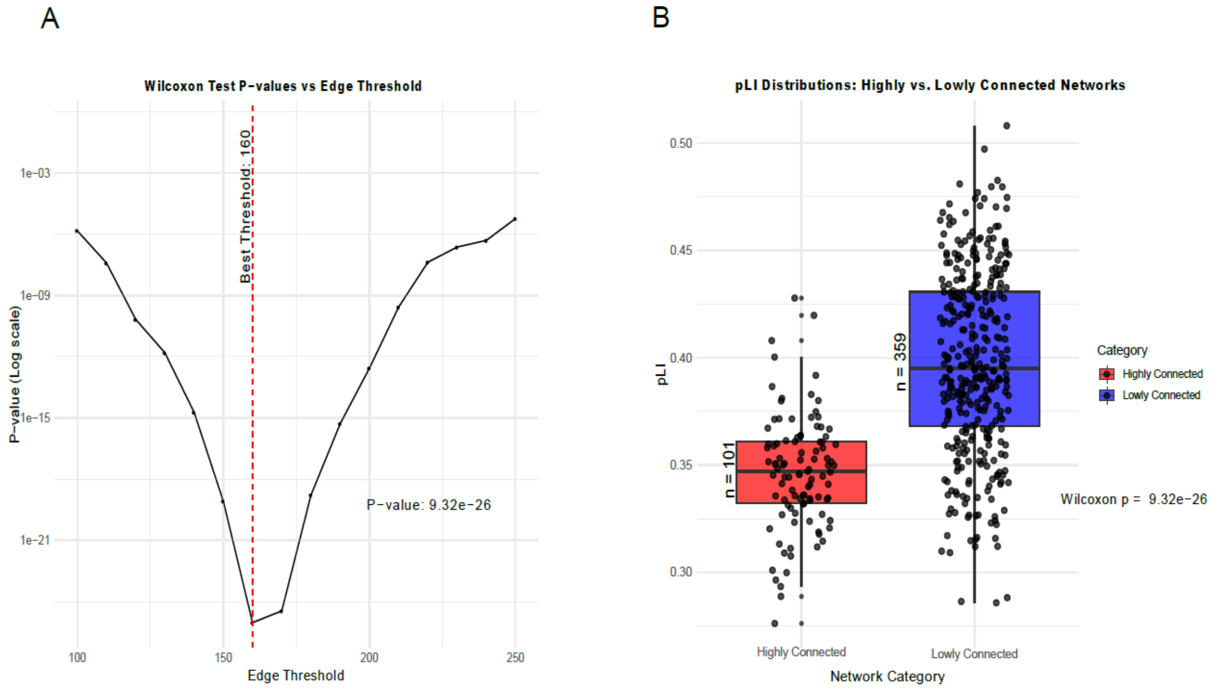


**Figure S7. Statistical Analysis of Network Connectivity and Gene Constraint Metrics.** A) Wilcoxon test p-values as a function of edge threshold. The statistical significance of differences in gene constraint metrics between highly connected and lowly connected networks is assessed using the Wilcoxon rank-sum test across various edge thresholds. The y-axis represents the p-value on a logarithmic scale, while the x-axis represents different edge thresholds. A sharp decline in p-values is observed as the edge threshold increases, indicating a stronger statistical separation between network categories. The optimal threshold is identified at 160 edges, yielding a highly significant *p*-value of 9.32e−26. B) Comparison of pLI score distributions for highly connected and lowly connected networks. The probability of being loss-of-function intolerant (pLI) is compared between two categories of networks: highly connected (red) and lowly connected (blue). The pLI score is a measure of gene essentiality, with higher values indicating genes that are more intolerant to loss-of-function mutations. The observed distribution suggests that genes in highly connected networks tend to have higher pLI scores compared to those in lowly connected networks, reinforcing the hypothesis that network centrality is correlated with gene constraint.

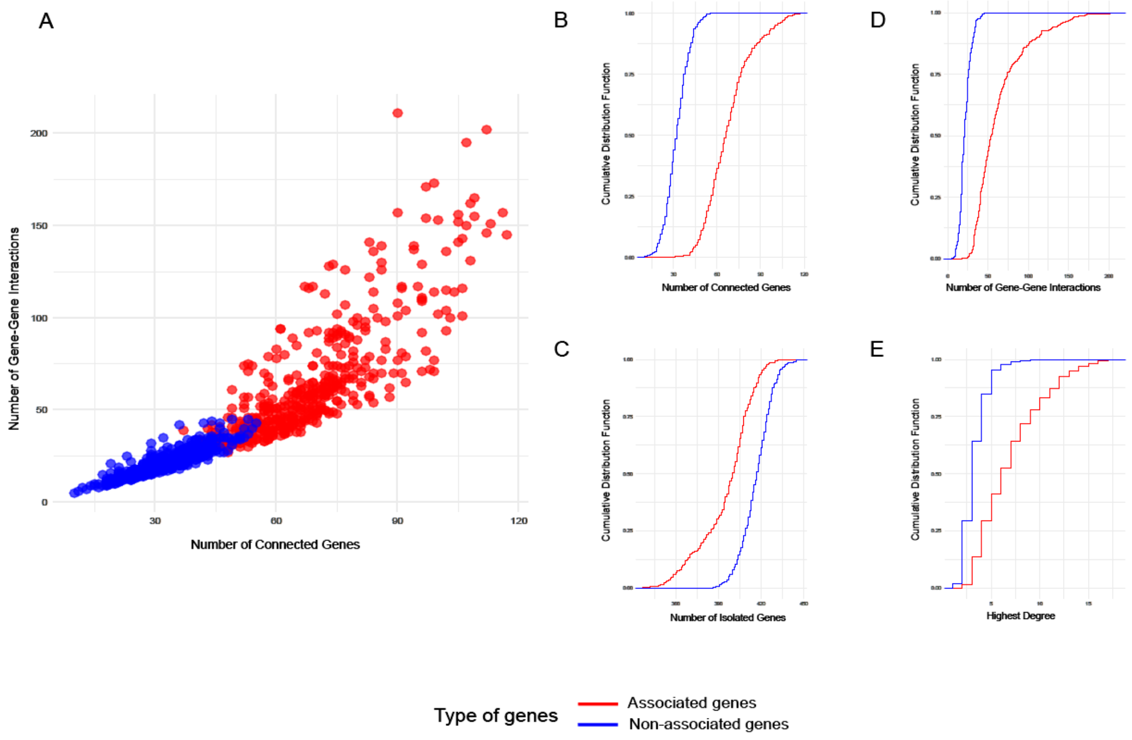


**Figure S8**. The comparison of gene-interaction networks built from physical interactions. A) The number of gene-gene interactions versus the number of connected genes in the interaction networks of associated genes (shown in red) and non-associated genes (shown in blue) with human polygenic phenotypes. Panels (B-E) present the cumulative distribution function (CDF) plots that compare various gene properties between associated and non-associated genes. In each plot, the y-axis represents the cumulative distribution, and the x-axis indicates the respective metric values: the number of connected genes (panel B), the number of isolated genes (panel C), the number of gene-gene interactions (panel D), and the highest degree of each network (panel D). The networks of associated genes (in red) and non-associated genes (in blue) were built using 500 highly, and 500 lowly associated genes, respectively. We selected the highly and lowly associated genes to each trait of interest from the MAGMA p-values. Comparisons of the network properties between associated and non-associated genes are highly significant, with $p<10^{-16}$ using the Wilcoxon rank-sum test.

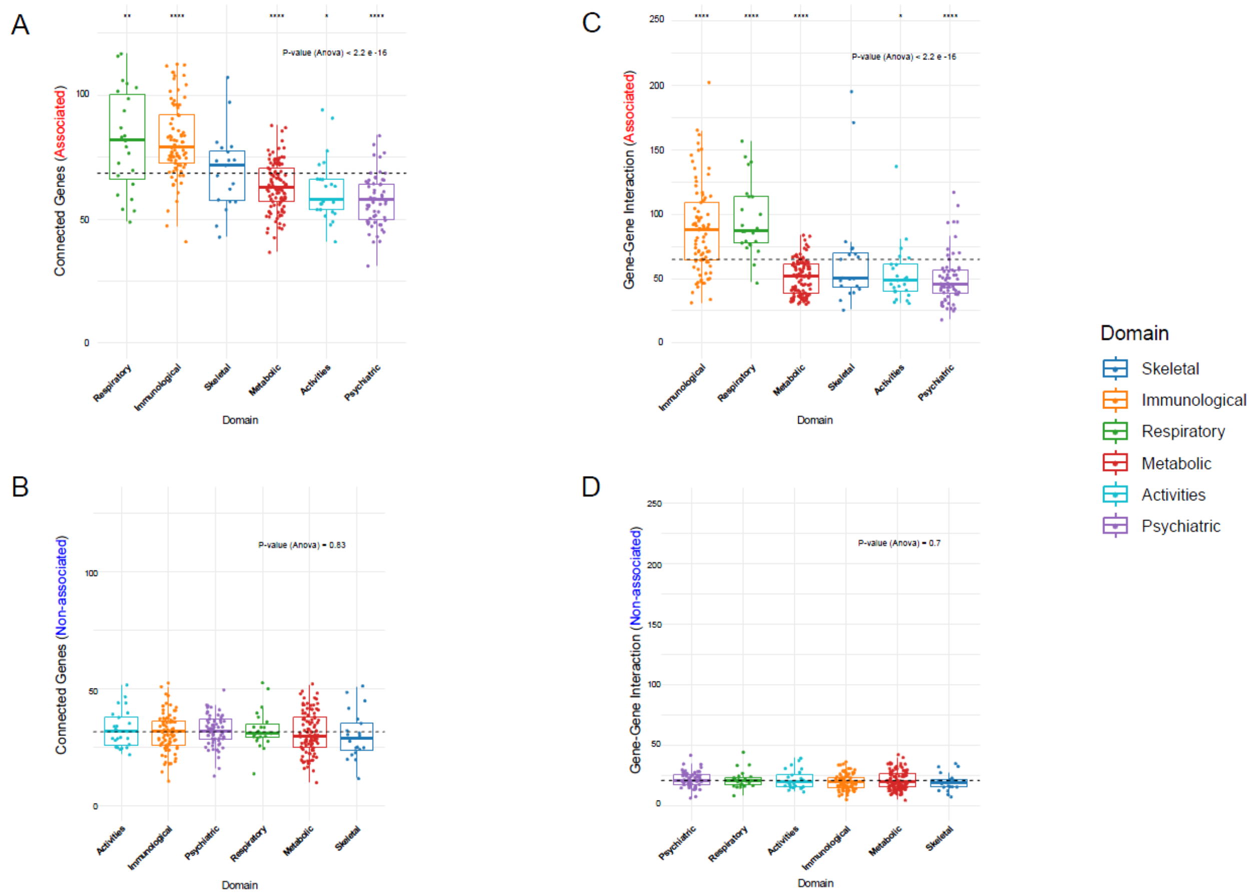


**Figure S9.** The connectivity of gene-interaction networks built from physical interactions varies across different domains of polygenic phenotypes. Comparative analysis of gene-gene interactions and connected genes across different biological domains for associated and non-associated genes. Panels (A-D) represent the distribution of gene interactions (A and B) and connectivity (C and D). (A) Gene-gene interactions in associated genes across biological domains including Skeletal, Immunological, Respiratory, Metabolic, Activities, and Psychiatric. (B) Gene-gene interactions in non-associated genes across the same domains. (C) Number of connected genes in associated genes and (D) in non-associated genes across the biological domains. Statistical significance is indicated by ANOVA p-values, with strong significance observed in Panels A and C (p-value < 2.2e-16), while B and D show non-significant variations (p-values of 0.83 and 0.7, respectively).

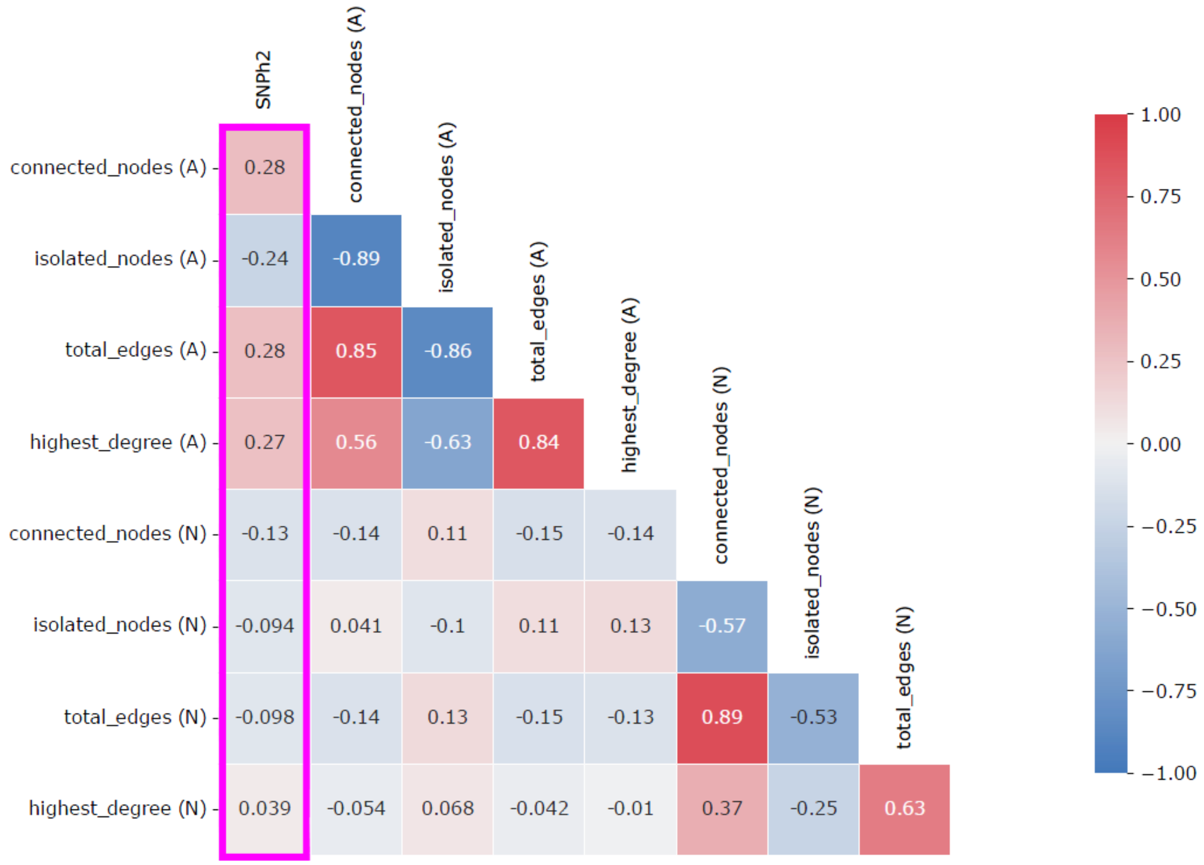


**Figure S10**. The correlation diagram showing the Spearman's rank correlation between SNP heritability of different polygenic traits and various properties of their gene interaction networks built from physical interactions.

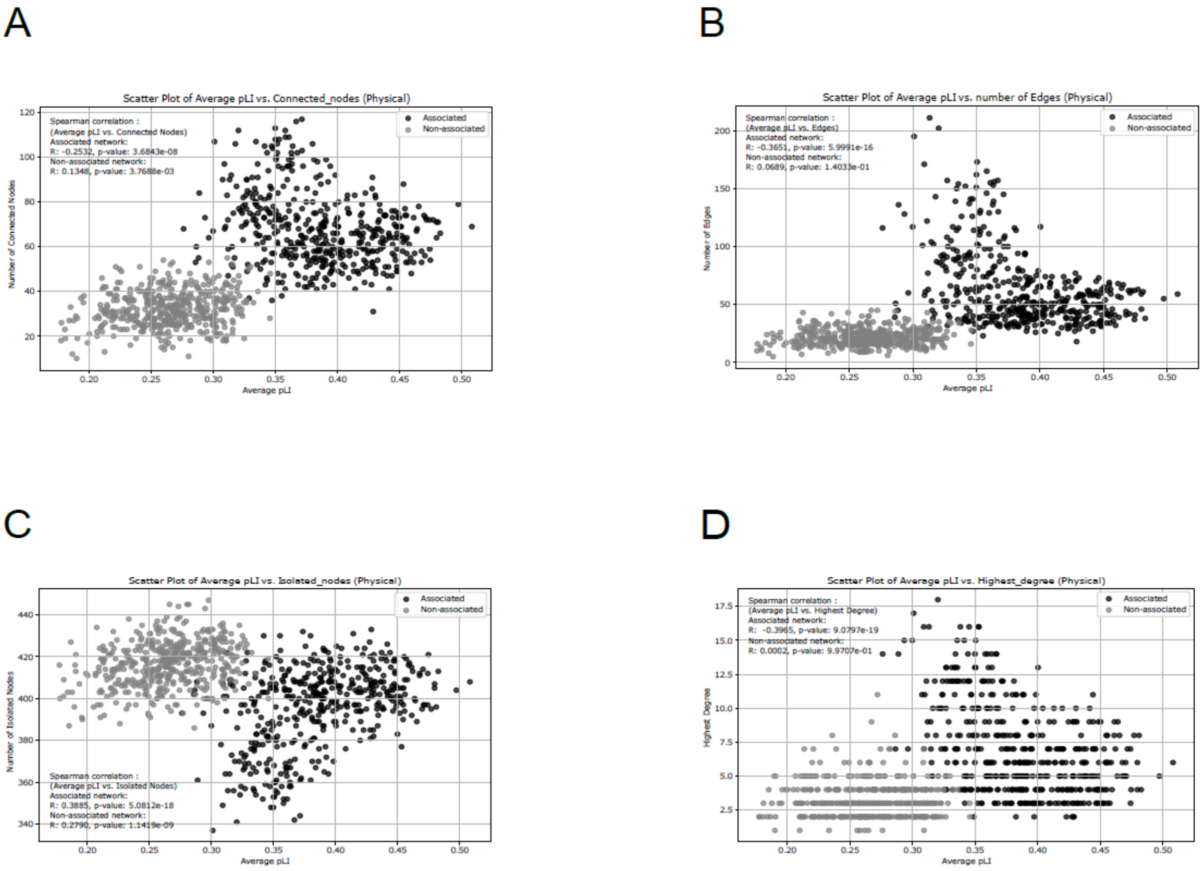


**Figure S11. Network characteristics and their correlation with pLI for gene interaction networks built from physical interactions.** A) Average pLI vs. Number of Connected Nodes: In the associated network, a weak negative correlation (R = -0.2532, p = 3.68e-08) suggests that highly connected genes tend to have lower pLI scores. No significant trend is observed in the non-associated network (R = 0.1348, p = 0.0037). (B) Average pLI vs. Number of Edges: A moderate negative correlation is observed in the associated network (R = -0.3651, p = 5.99e-16), while the non-associated network shows no significant trend (R = 0.0689, p = 0.14). (C) Average pLI vs. Number of Isolated Nodes: Both networks show a positive correlation, with isolated genes tending to have higher pLI values (associated: R = 0.3885, p = 5.08e-18; non-associated: R = 0.2790, p = 1.14e-09). (D) Average pLI vs. Highest Degree: A negative correlation suggests that highly connected genes tend to have lower pLI scores (associated: R = -0.3965, p = 9.07e-19; non-associated: R = 0.0002, p = 0.997). These findings suggest that in the associated network, higher connectivity is linked to lower gene constraint, while isolated genes exhibit greater functional constraint. This trend is weaker in the non-associated network.

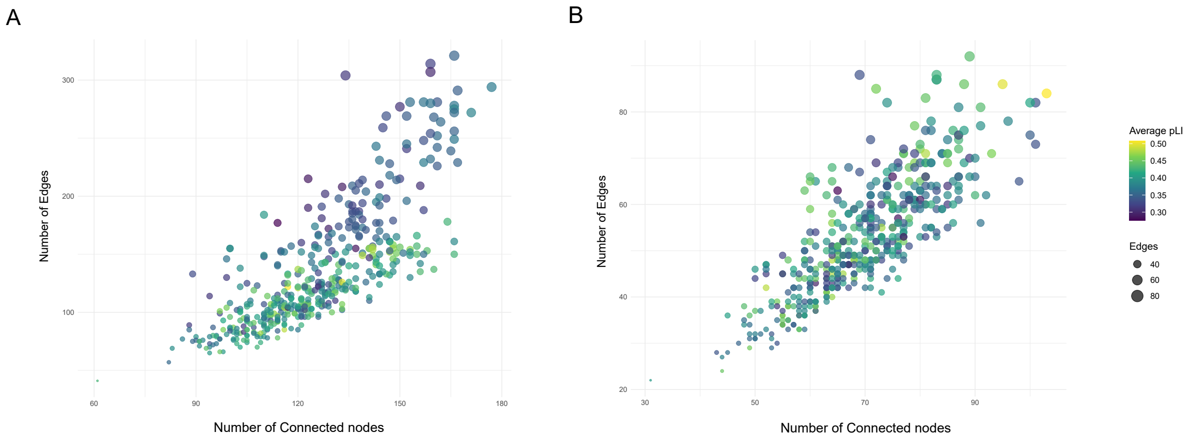


**Figure S12. Network architecture of 461 human polygenic phenotypes built from functional interaction scores in relation to the average pLI of the embedded genes**. A) The number of edges in each network versus the number of connected nodes for the gene interaction networks of the associated genes. B) The number of edges in each network versus the number of connected nodes for the gene interaction networks of the non-associated genes. In both panels the size of each point (phenotype) is proportional to the number of edges (y-axis values), and the color intensity changes from blue (low average pLI) to green (high average pLI).

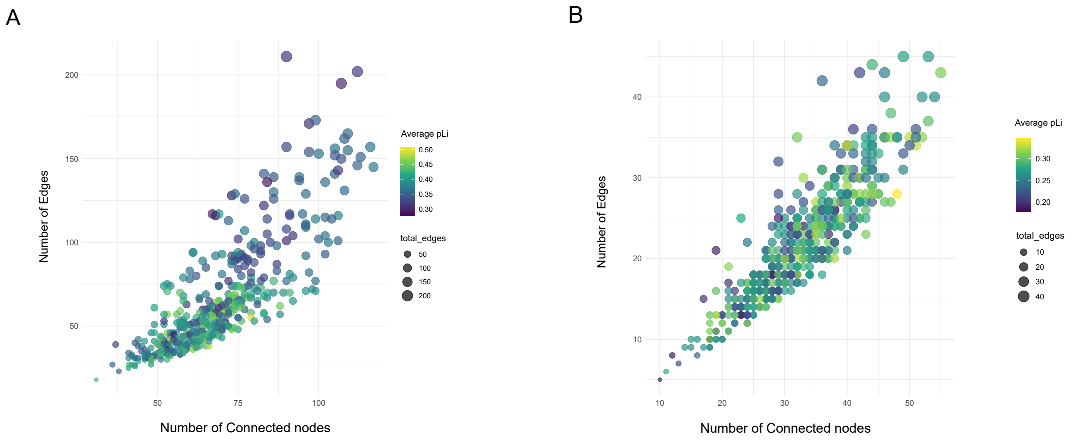


**Figure S13**. **Network architecture of 461 human polygenic phenotypes built from physical interaction scores in relation to the average pLI of the embedded genes**. A) The number of edges in each network versus the number of connected nodes for the gene interaction networks of the associated genes. B) The number of edges in each network versus the number of connected nodes for the gene interaction networks of the non-associated genes. In both panels the size of each point (phenotype) is proportional to the number of edges (y-axis values), and the color intensity changes from blue (low average pLI) to green (high average pLI).

### Supplementary Note 1

We performed multiple linear correlation between SNPh2 of all 4756 human polygenic phenotypes in the GWAS atlas, GWAS power (N=number of participants), and the number of edges of the gene interaction networks of the associated genes built from the functional and physical scores (Tables S8 and S9).

*Functional networks*

```
Call:
lm(formula = SNPh2 ~ ., data = df)

Residuals:
     Min       1Q   Median       3Q      Max 
-0.58329 -0.06397 -0.01960  0.04553  0.99733 

Coefficients:
              Estimate Std. Error t value Pr(>|t|)    
(Intercept)  9.755e-02  6.792e-03  14.363  < 2e-16 ***
Edges        5.962e-04  7.496e-05   7.953 2.85e-15 ***
N           -2.573e-07  1.708e-08 -15.066  < 2e-16 ***
---
Signif. codes:  0 ‘***’ 0.001 ‘**’ 0.01 ‘*’ 0.05 ‘.’ 0.1 ‘ ’ 1

Residual standard error: 0.1238 on 2258 degrees of freedom
  (2495 observations deleted due to missingness)
Multiple R-squared:  0.09974,	Adjusted R-squared:  0.09894 
F-statistic: 125.1 on 2 and 2258 DF,  p-value: < 2.2e-16
```

*Physical networks*

```
Call:
lm(formula = SNPh2 ~ ., data = df)

Residuals:
     Min       1Q   Median       3Q      Max 
-0.59039 -0.06523 -0.02069  0.04425  0.99121 

Coefficients:
              Estimate Std. Error t value Pr(>|t|)    
(Intercept)  1.102e-01  5.564e-03   19.81  < 2e-16 ***
Edges        9.643e-04  1.249e-04    7.72 1.73e-14 ***
N           -2.538e-07  1.702e-08  -14.91  < 2e-16 ***
---
Signif. codes:  0 ‘***’ 0.001 ‘**’ 0.01 ‘*’ 0.05 ‘.’ 0.1 ‘ ’ 1

Residual standard error: 0.1239 on 2258 degrees of freedom
  (2495 observations deleted due to missingness)
Multiple R-squared:  0.09832,	Adjusted R-squared:  0.09752 
F-statistic: 123.1 on 2 and 2258 DF,  p-value: < 2.2e-16
```

**Supplementary Note 2**

Since expression level is a key determinant of evolutionary rate, we accounted for this factor by calculating the partial correlation between average d*N*/d*S* and various network properties, such as the number of edges, connected nodes, isolated nodes, and the highest degree, for both associated and non-associated networks. The correlations between network properties and d*N*/d*S* remained significant for associated networks even after controlling for expression level ($R$ = 0.052, $p$ = $1.44 \times 10^{-7}$, Spearman's rank correlation), whereas no significant correlations were observed for non-associated networks ($R$ = -0.0029, $p$ = 0.76, Spearman's rank correlation).
